# Modelling the coupling of the M-clock and C-clock in lymphatic muscle cells

**DOI:** 10.1101/2021.11.07.467629

**Authors:** E.J. Hancock, S.D. Zawieja, C. Macaskill, M.J. Davis, C.D. Bertram

## Abstract

Lymphoedema develops due to chronic dysfunction of the lymphatic vascular system which results in fluid accumulation between cells. The condition is commonly acquired secondary to diseases such as cancer or the therapies associated with it. The primary driving force for fluid return through the lymphatic vasculature is provided by contractions of the muscularized lymphatic collecting vessels, driven by electrical oscillations. However, there is an incomplete understanding of the molecular and bioelectric mechanisms involved in lymphatic muscle cell excitation, hampering the development and use of pharmacological therapies. Modelling *in silico* has contributed greatly to understanding the contributions of specific ion channels to the cardiac action potential, but modelling of these processes in lymphatic muscle remains limited. Here, we propose a model of oscillations in the membrane voltage (M-clock) and intracellular calcium concentrations (C-clock) of lymphatic muscle cells. We modify a model by Imtiaz and colleagues to enable the M-clock to drive the C-clock oscillations. This approach differs from typical models of calcium oscillators in lymphatic and related cell types, but is required to fit recent experimental data. We include an additional voltage dependence in the gating variable control for the L-type calcium channel, enabling the M-clock to oscillate independently of the C-clock. We use phase-plane analysis to show that these M-clock oscillations are qualitatively similar to those of a generalised FitzHugh-Nagumo model. We also provide phase plane analysis to understand the interaction of the M-clock and C-clock oscillations. The model and methods have the potential to help determine mechanisms and find targets for pharmacological treatment of lymphoedema.

## 1 INTRODUCTION

The lymphatic vascular system returns interstitial (extracellular) fluid from all over the body to the subclavian veins, thereby maintaining homeostasis in the face of necessary efflux of fluid from blood capillaries. In the process the system also transports viruses, bacteria and particulate matter to lymph nodes where they are mechanically filtered and/or biologically neutralized by immune cells. The advection of antigen from the body’s cells to lymph nodes, either in solution or carried by antigen presenting cells, provides rapid notification to the adaptive immune system, allowing a prompt whole-body response targeted at the specific antigen. Lymphatic vascular defects are now known to be involved in a wide range of diseases, including obesity, cardiovascular disease, inflammation, hypertension, atherosclerosis, Crohn’s disease, glaucoma and neurological disorders such as Alzheimer’s disease (Oliver et al. 2020).

Unlike blood flow, lymph flow is not propelled by the heart but relies on the intrinsic contractile activity of the muscular collecting vessels. The lymphatic vascular system consists of three functional types of vessel: initial, pre-collecting and collecting. Interstitial fluid becomes lymph when it enters the porous initial lymphatics, entry being favoured by a specialized arrangement of endothelial cells forming primary valves (Trzewik et al. 2001, Baluk et al. 2007). Lymph flow within the lymphatic vascular system is the result of two types of pumping, both dependent on the subdivision of collecting vessels into short segments by one-way secondary valves; less frequent secondary valves also occur in pre-collectors (Breslin et al. 2019). Intrinsic pumping, which accounts for two-thirds of lymph flow at rest in the extremities (Engeset et al. 1977, Olszewski & Engeset 1980), then results from the rhythmic contraction of muscle in the wall of the collectors, causing each segment to expel fluid in the direction prescribed by the valves. Extrinsic pumping is the result of passive squeezing of the valved vessels through relative motion of surrounding tissues.

When lymphatic fluid is not effectively transported, as a result of dysfunction of the lymphatic vascular system, the ensuing condition, in which interstitial fluid accumulates in one or more body regions, is called lymphoedema. It consists of sustained swelling in the affected region, aberrant fibrosis, adipose accumulation and disruption of the interstitial composition, and chronic lymphoedema predisposes to secondary infections. The dysfunction can be inherited (primary lymphoedema) but is commonly acquired secondary to diseases such as cancer and congestive heart failure. Lymphoedema affects over 10 million people annually in the USA (Rockson & Rivera 2008), and over 130 million people worldwide (Mortimer & Rockson 2014). Despite its prevalence, there are currently no pharmacological treatments and there is an incomplete understanding of the mechanisms involved. Current therapies utilise external massage and compression garments for temporary relief of symptoms (Adams et al. 2010), but do not address the underlying causes. Lymphoedema can be the result of either incompetent secondary valves, inadequate lymphatic muscle function, or overload of the lymphatic pump (Scallan et al. 2016). Overall, lack of active lymph pumping, dependent on the coordinated actions of lymphatic muscle and valves, is central to this disease entity.

Our present understanding of the underlying mechanisms leading to lymphatic contractile dysfunction is primitive. Lymphatic muscle is uniquely complex (Muthuchamy et al. 2003), in that it combines a fast phenotype capable of maintaining rhythmic twitch contractions indefinitely, a slow phenotype responsible for sustained lymphatic vascular tone, and an inbuilt pace-maker for the twitch contractions. In addition, it mounts sensitive inotropic and chronotropic responses to variations in local lymphatic distending pressure and lymph flow-rate (Gashev 2008). A much better understanding of the molecular and ionic mechanisms regulating lymphatic muscle is needed, and this demands both detailed examination of lymphatic muscle cell (LMC) behaviour in the biology laboratory and parallel modelling of the complex cellular systems that provide these multiple functions, including both autonomous pace-making and coordination of the contractions of adjacent collecting lymphatic segments.

The classical models of pace-making in cardiac cells (Noble 1962, McAllister et al. 1975) have more recently evolved to a paradigm based on the coupled interactions of a so-called ‘membrane clock’ (M-clock) and a ‘calcium clock’ (C-clock) (Maltsev & Lakatta 2013, Yaniv et al. 2015). The M-clock comprises all the equations dealing with ion passage through the cell membrane and the resulting voltage oscillations. The C-clock focuses on the cyclical release of Ca^2+^ into the cytoplasm from intracellular stores, via receptors sensitive to either inositol trisphosphate (IP3) or ryanodine.

Another cell type which incorporates an autonomous pace-maker is the interstitial cell of Cajal (ICC), which activates gastrointestinal smooth muscle to produce peristaltic transport of gut contents (Sanders 2019). Although there are important differences between the dominant ion channels in ICCs and in LMCs (To et al. 2020), and ICCs do not house the actual calcium-regulated contractile filaments as do LMCs, this is otherwise considered the closest cell type to LMCs in respect of its pace-making, producing rhythmic contractions at comparably slow rates. Complex models of coupled calcium and voltage oscillations in ICCs have been developed (Lees-Green et al. 2014, Youm et al. 2019). In particular, these ICC models highlight the role of the calcium-activated anoctamin-1 (Ano1) chloride-ion channel in ICC oscillations; this channel is known to play an important role in LMCs as well (Zawieja et al. 2019). However, these models are primarily based around the C-clock; if the Ano1 channel is disabled, voltage oscillations become small or non-existent, while large-scale [Ca^2+^] oscillations continue. Furthermore, the detailed mechanistic nature of these models (the model of Youm et al. consists of 23 ordinary differential equations) presents difficulties in qualitatively analysing and interpreting C-clock and M-clock interactions, and requires information yet to be acquired in lymphatic muscle.

The arrangement of the ICC models, whereby the M-clock is effectively enslaved to a master C-clock, does not accord with our recent experimental data from LMCs, where voltage oscillations continue after knockout of Ano1 channels; see Figure 1. Although the frequency of action potentials reduces by a factor of about four after Ano1 knock-out, the action potential itself is maintained, and even increases in both maximum depolarized voltage and total peak-to-peak voltage excursion.

**Figure 1.**
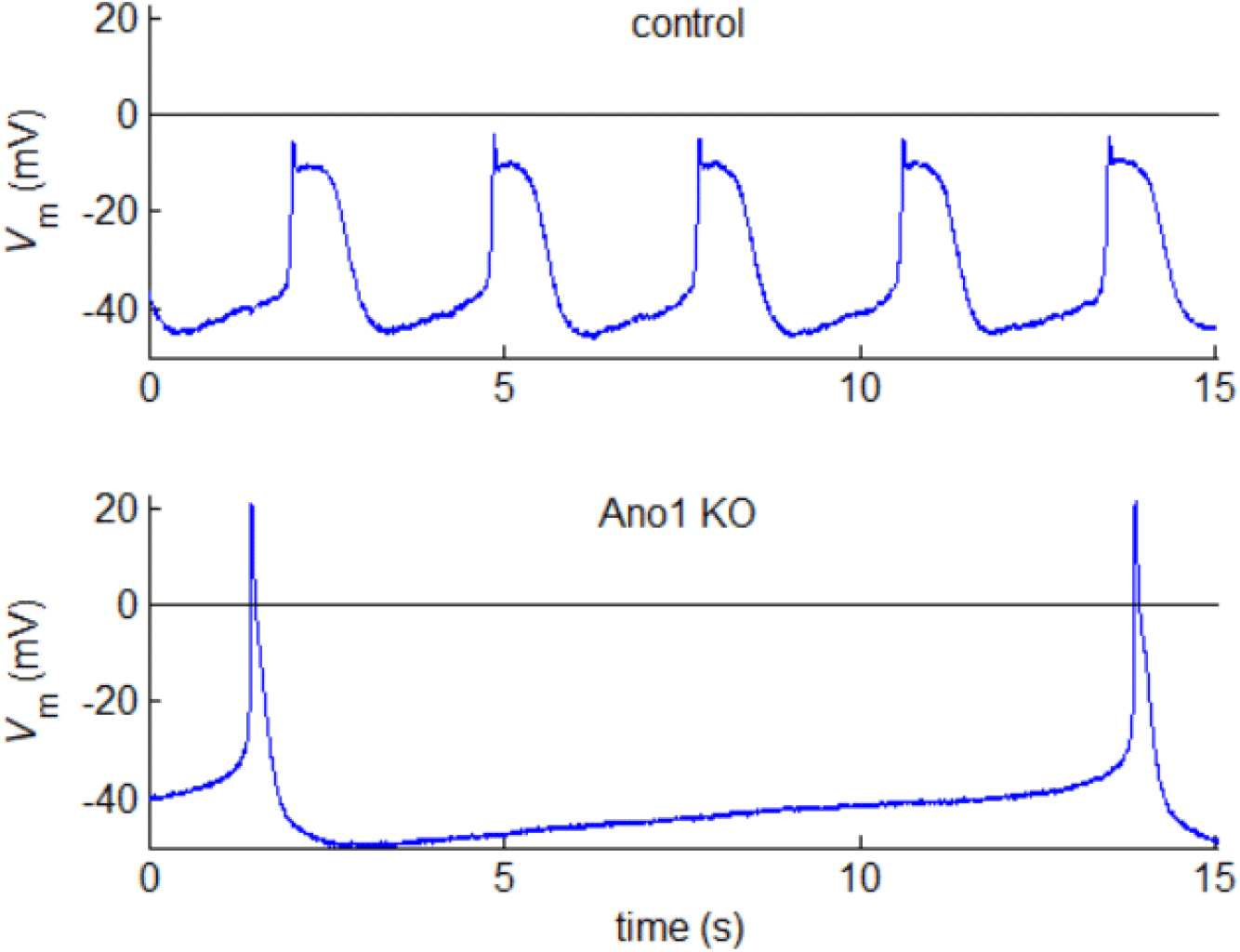
Membrane voltage recordings from single muscle cells in murine inguinal axillary lymphatic vessels. In this case, Ano1-KO was the result of genetic deletion (inducible smooth-muscle-specific Ano1 KO mouse).

In contrast to other cell types, the current modelling of oscillations in LMCs is limited. Imtiaz et al. (2007) added an M-clock component to an earlier calcium-based model by Dupont & Goldbeter (1993). Similarly, the model of Hald et al. (2018) is based on the two-variable voltage oscillator of Morris & Lecar (1981), which posits an excitable system of two nonlinear conductances, and was developed to explain the behaviour of barnacle muscle. In the Imtiaz model, as in the ICC models, the store-controlled calcium oscillator can function independently of whether action potentials occur, at a frequency which is essentially unchanged by disabling calcium-activated chloride exit from the cell. Since the current in question is of chloride ions, the result of Ano1 channels opening is to change the membrane voltage, thereby affecting voltage-gated ion channels on the cell membrane and thus the action of the M-clock, but there is no direct effect on calcium concentrations, i.e., the C-clock. Again, this behaviour is contrary to the observations shown in Figure 1.

Here, we propose a model of oscillations in lymphatic muscle cells whereby the M-clock drives the C-clock, enabling the model to match the data in Figure 1. The proposed model modifies the existing model of Imtiaz et al. (2007) by the inclusion of an additional voltage dependence in the gating variable control for the L-type calcium channel. L-type current is essential for lymphatic pace-making, action potentials and contractions (To et al. 2020), whereas it is not the essential calcium channel in sino-atrial node and ICCs. We first analyse the existing model, showing how its simulations are inconsistent with the data. We use phase-plane analysis to comprehend the underlying reasons, providing a qualitative understanding of the C-clock oscillations and how they drive the M-clock oscillations. We then go on to use phase-plane analysis and waveform simulations to study the proposed model, showing its consistency with the data in Figure 1. We show that this model’s M-clock is qualitatively similar to a generalised FitzHugh-Nagumo model (FitzHugh 1961, Nagumo et al. 1962, Keener & Sneyd 2009), enabling it to oscillate independently. We also introduce phase-plane analysis of the interaction of the M-clock and C-clock oscillations, illustrating how the C-clock and Ano1 modify the frequency of M-clock oscillations in the new model.

The remainder of the paper is set out as follows. Section 2 describes the methods used in the animal experiments and in the computer models. Section 3 presents the results from first our reworked version of the existing model from Imtiaz et al. (2007), which will be referred to henceforth as the existing model, and then our new M-clock-centred model. Section 4 discusses these results, along with more general issues relating to modelling and the extent to which our modifications are in line with experimental evidence. A short Section 5 then restates the main results in the form of conclusions.

## 2 METHODS

### 2.1 Animal experiments

The experimental data were obtained in experiments on isolated segments of mouse inguinal-axillary lymphatic vessel. A detailed description of the methods employed is already available (Zawieja et al. 2019), so only a brief summary will be given here. All animal protocols were approved by the institutional Animal Care and Use Committee at the University of Missouri, and in accordance with the NIH Guide for the Care and Use of Laboratory Animals.

Inguinal-axillary lymphatic vessels were dissected from Nembutal-anaesthetized male mice. Half-lymphangion segments of vessel were cleaned of adherent external tissue and mounted on glass micropipettes of 80 μm diameter for pressurisation and subsequent sharp electrode impalement, all whilst immersed in physiological saline solution. The preparation was moved to the stage of an inverting microscope, from which vessel images could be recorded continuously. With the pipettes connected to reservoirs supplying an identical pressure of 8 cmH_2_O upstream and downstream, the distance between micropipettes was adjusted such that vessel buckling was avoided, then the pressure was reduced to 3 cmH_2_O. To reduce vessel wall movement through spontaneous contractile activity while leaving the underlying electrical activity unaffected, 2 μM wortmannin, which paralyses by inhibiting myosin light chain kinase, was added to the perfusion bath, and a period of 30–60 mins was allowed for the contractions to reach < 5 μm diameter reduction as monitored through video tracking by a custom algorithm (Davis 2005). Lymphatic muscle cell (LMC) impalement was by microelectrode filled with 1 M KCl having a resistance of 250–350 MΩ. Membrane voltage was sampled at 1 kHz or greater and recorded using a custom Labview programme. For pharmacological inhibition of Ano1, benzbromarone was added to the bath to produce a concentration of 5 μM. For genetic deletion of Ano1, inguinal-axillary vessels from Ano1^ismKO^ mice were used; see Zawieja et al. (2019) for details of the induction protocols. Since benzbromarone can have off-target effects, and genetic deletion may not be total, further experiments (unpublished data) used the highly specific Ano1 inhibitor Ani9 (Seo et al. 2016) at 3 μM concentration. All three methods gave similar results.

### 2.2 Electrochemical model

In this section we introduce a model for LMC oscillations. The model is modified from a LMC model by Imtiaz et al. (2007), itself based around a model of calcium oscillations alone by Dupont & Goldbeter (1993). The model incorporates both the M-clock and C-clock oscillations using four variables; see Figure 2. The C-clock is represented by two calcium concentration variables: the cytosolic calcium concentration *Z* and the ER (endoplasmic reticulum) stored calcium concentration *Y*. The M-clock is represented by two membrane variables: the membrane potential *V* and an inactivation gating variable *h* for the L-type calcium channel. The C-clock is dependent on the fluxes through the membrane calcium channels, including the L-type channel, and the calcium channels between the cytosol and the intracellular calcium store (ER). The membrane potential varies with the ion fluxes through the membrane. The model is represented by

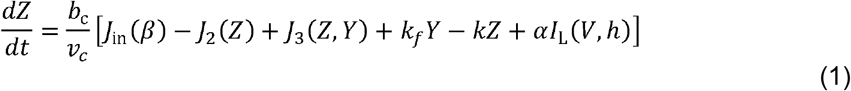

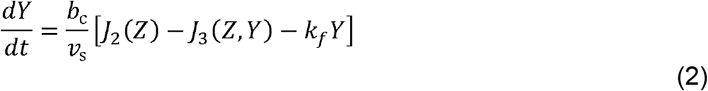

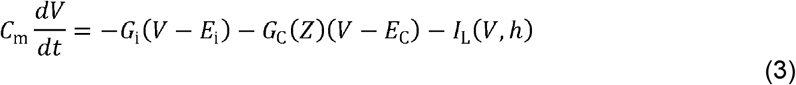

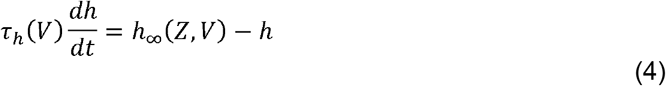

where *J*_in_ is the Ca^2+^ influx across the membrane into the cell from outside, *β* is the cytosolic concentration of IP3 (herein chosen to remain constant), *J*_2_ is the Ca^2+^ influx from the cytosol into the store, *J*_3_ is the Ca^2+^-dependent and IP3-dependent release of calcium from the store into the cytosol, *k*_f_*Y* represents leakage from the store and *kZ* represents the pumping of Ca^2+^ from the cytosol out of the cell. For the membrane voltage, *C*_m_ represents the capacitance of the cell membrane, *G*_i_ and *E*_i_ represent a lumped conductance and equilibrium potential respectively for passive non-selective ionic channels, *G*_C_ and *E*_C_ represent a lumped conductance and equilibrium potential respectively for the Ano1 channel, and *I*_L_ represents the current in the L-type calcium channel. *h*_∞_ and *τ_h_* represent the steady state and voltage-dependent time constant respectively for the L-type inactivation gating variable *h*. *v*_c_ is the volume of the cytoplasm, *v*_s_ is the volume of the ER, *b*_c_ represents the fraction of total calcium that is unbuffered, and *α* is a conversion factor between current and calcium ion flux. These functions are described by

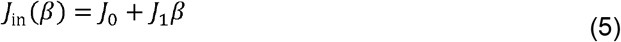

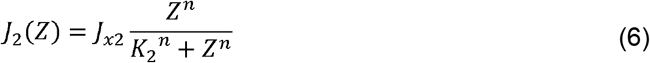

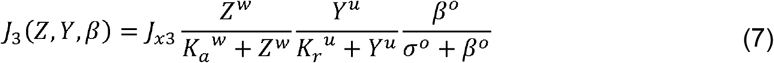

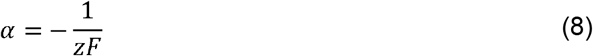

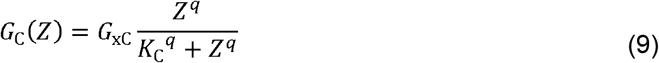

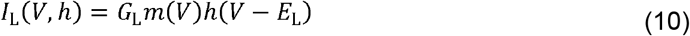

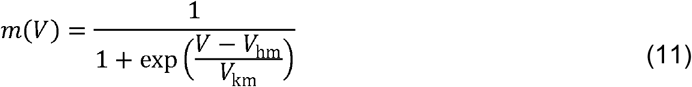

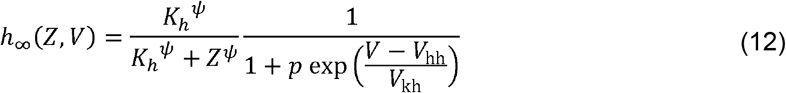

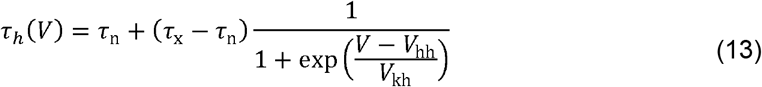

where the baseline values of all constants are given in Table 1. The L-type current is controlled by the activating and inactivating gate variables, *m* and *h* respectively, where *m* is assumed to act instantaneously and *h* is subject to first-order delay kinetics according to the time constant *τ_h_*, itself now ranging between maximum and minimum values *τ*_x_ and *τ*_n_ respectively according to the membrane voltage.

**Figure 2.**
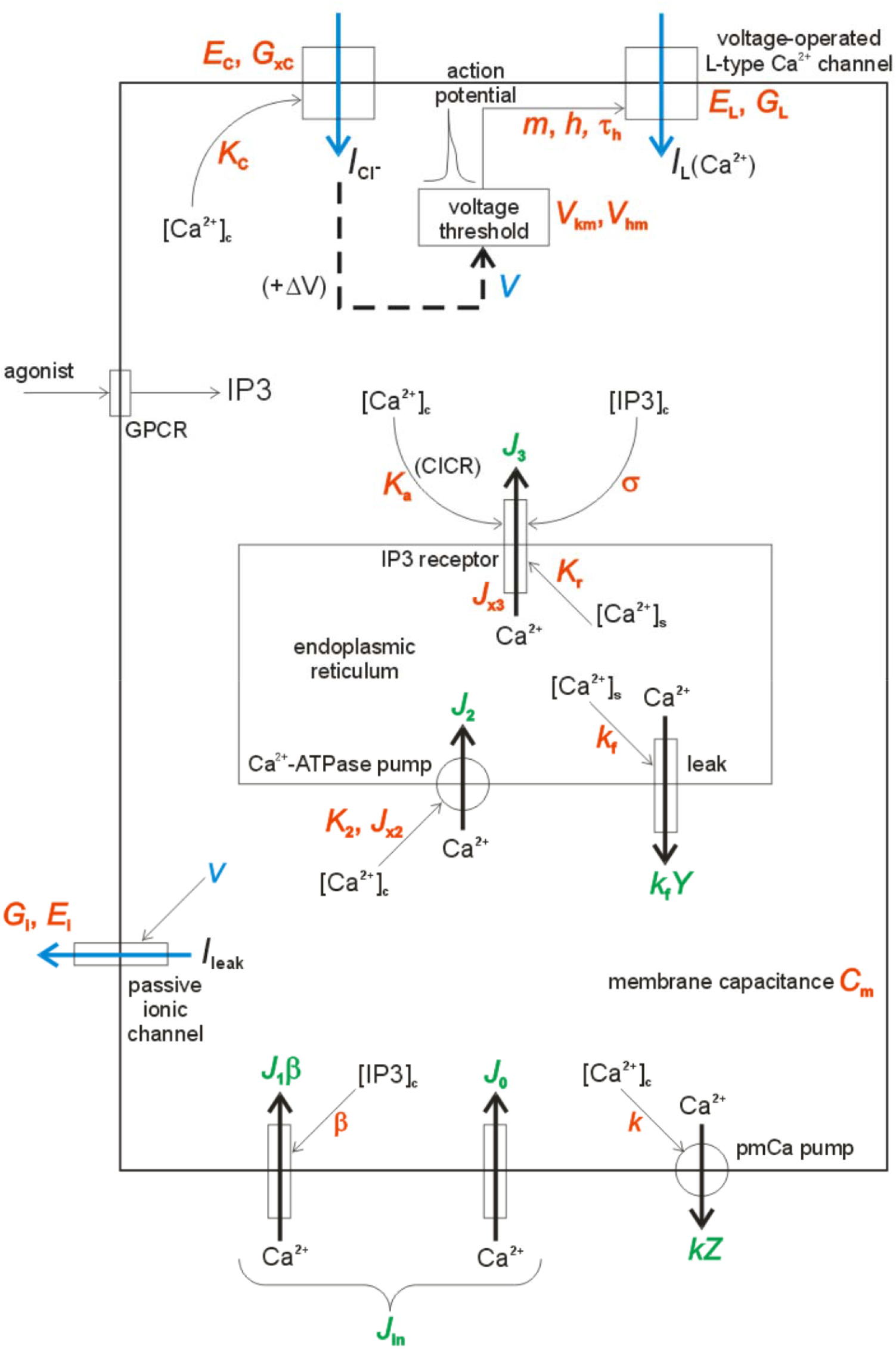
Schematic of the LMC model. Calcium fluxes are shown by black arrows; other currents, plus the L-type calcium current (blue arrows) are shown in the direction of predominant motion of positive charges, irrespective of the actual ion. CICR = calcium-induced calcium release, GPCR = G-protein-coupled receptor. Suffix c = cytosol, s = store (endoplasmic reticulum). *V* = voltage, *Y* = store [Ca^2+^], *Z* = cytosolic [Ca^2+^]. See equations and Table 1 for the meaning of all other symbols.

**Table 1.** Baseline values for constants in the existing model, and as modified for the new model. In the source column, 1 denotes Imtiaz et al. (2007), although in many cases the values therein have been reinterpreted in terms of units, etc., and 2 denotes Lees-Green et al. (2014).

| parameter | units | meaning | existing value | source | new value |
| --- | --- | --- | --- | --- | --- |
| $\theta$ | $\mu\text{M}$ | cytosolic [IP3] | 0.4 | 1 | |
| $J_0$ | $\mu\text{mol/s}$ | [IP3]-independent $\text{Ca}^{2+}$ influx | $5.67 \times 10^{-12}$ | 1 | |
| $J_1$ | $(\mu\text{mol/s})/\mu\text{M}$ | cytosol $\text{Ca}^{2+}$ influx per unit [IP3] | $5.67 \times 10^{-12}$ | 1 | |
| $k$ | /s | rate constant, $\text{Ca}^{2+}$ pumping from cell | 16.7 | 1 | |
| $J_{x2}$ | $\mu\text{mol/s}$ | max. $\text{Ca}^{2+}$ pumping rate into store | $8.33 \times 10^{-11}$ | 1 | |
| $n$ | - | $J_2(Z)$ Hill coeft. | 2 | 1 | |
| $K_2$ | $\mu\text{M}$ | $[\text{Ca}^{2+}]_c$-dependent threshold for $\text{Ca}^{2+}$ flux into store | 1 | 1 | |
| $k_f$ | /s | rate constant for $\text{Ca}^{2+}$ leak from store | 1.67 | 1 | |
| $J_{x3}$ | $\mu\text{mol/s}$ | max. rate of $\text{Ca}^{2+}$ release from store | $1.0833 \times 10^{-9}$ | 1 | |
| $o$ | - | $J_3(\theta)$ Hill coeft. | 4 | 1 | |
| $u$ | - | $J_3(Y)$ Hill coeft. | 2 | 1 | |
| $w$ | - | $J_3(Z)$ Hill coeft. | 4 | 1 | |
| $K_r$ | $\mu\text{M}$ | $[\text{Ca}^{2+}]_s$-dependent threshold for store $\text{Ca}^{2+}$ release | 20 | 1 | |
| $K_a$ | $\mu\text{M}$ | $[\text{Ca}^{2+}]_c$-dependent threshold for store $\text{Ca}^{2+}$ release | 0.9 | 1 | |
| $\sigma$ | $\mu\text{M}$ | [IP3]-dependent threshold for store $\text{Ca}^{2+}$ release | 0.48 | 1 | |
| $K_C$ | $\mu\text{M}$ | half saturation for $\text{Ca}^{2+}$-induced $\text{Cl}^-$ conductance | 1.4 | 1 | 3 |
| $q$ | - | $G_C(Z)$ Hill coeft. | 4 | 1 | |
| $b_c$ | - | $\text{Ca}^{2+}$ buffering constant | 0.01 | 2 | |
| $v_c$ | L | cytosol volume | $10^{-12}$ | 2 | |
| $v_s$ | L | store volume | $10^{-13}$ | 2 | |
| $C_m$ | pF | cell capacitance | 25 | 2 | |
| $E_i$ | mV | reversal potential, passive channels | -67.2 | 1 | |
| $E_C$ | mV | reversal potential for Ano1 current | -20 | 1 | |
| $E_L$ | mV | reversal potential for L-type channels | +20 | 1 | |
| $G_i$ | nS | conductance of passive channels | 1.12 | 1 | |
| $G_{xC}$ | nS | max. $\text{Ca}^{2+}$-induced $\text{Cl}^-$ conductance | 4 | 1 | 0.25 |
| $G_L$ | nS | conductance of L-type $\text{Ca}^{2+}$ channels | 5 | 1 | |
| $V_{hm}$ | mV | half saturation for gate fn. $m(V)$ | -42 | 1 | -46 |
| $V_{km}$ | mV | inverse slope const., gate fn. $m(V)$ | -2 | 1 | -4 |
| $K_h$ | $\mu\text{M}$ | half saturation for gate fn. $h_\infty(Z)$ | 0.7 | 1 | 4 |
| $\psi$ | - | Hill coeft. for $h_\infty(Z)$ | 20 | 1 | |
| $V_{hh}$ | mV | half saturation for gate fn. $h_\infty(V)$ | -60 | | -48 |
| $V_{kh}$ | mV | inverse slope const., gate fn. $h_\infty(V)$ | 1 | | |
| $p$ | - | V-dependence of L-type inactivation | 0 | | 1 |
| $\tau_n$ | s | minimum of time constant $\tau_h(V)$ | 1.2 | 1 | 0.5 |
| $\tau_x$ | s | maximum of time constant $\tau_h(V)$ | 1.2 | 1 | 10 |
| $\alpha$ | ($\mu\text{mol/s}$)/pA | $\text{Ca}^{2+}$ flux per unit current | $-5.18 \times 10^{-12}$ | | |
| $z$ | - | valency of calcium ion | 2 | | |
| $F$ | pC/$\mu\text{mol}$ | Faraday constant | $9.6485 \times 10^{10}$ | | |

## 3 RESULTS

In this section we present waveforms from and phase plane analysis for the existing and proposed models of oscillations in membrane potential and calcium concentration. We first show that the existing model, based on that of Imtiaz et al. (2007), does not qualitatively replicate the experimental Ano1-KO data for LMCs, before showing that the proposed model with modifications to the L-type channel model qualitatively matches the data.

### 3.1 Analysis of the existing model

We first created simulations of lymphatic pace-making by integrating the equations of the existing model, with and without the inclusion of the Ano1 channel, to produce the results in Figure 3. For this existing case, the steady-state inactivation gating variable *h*_∞_ depends only on the cytosolic calcium concentration *Z*(*t*); there is no direct voltage dependence of the inactivation for the L-type calcium channel. This is achieved by setting *p* = 0 in eq. 12 and *τ_x_* = *τ_n_* in eq. 13. In Figure 3, the wild-type system which includes the Ano1 channel produces regular oscillations which include well-formed action potentials (APs), but Ano1 knock-out (by setting *G*_xC_ = 0 in eq. 9) eliminates these oscillations. This observation contrasts with the experimental data shown in Figure 1, where, after Ano1 knock-out, oscillations continue but at a lower frequency.

**Figure 3.**
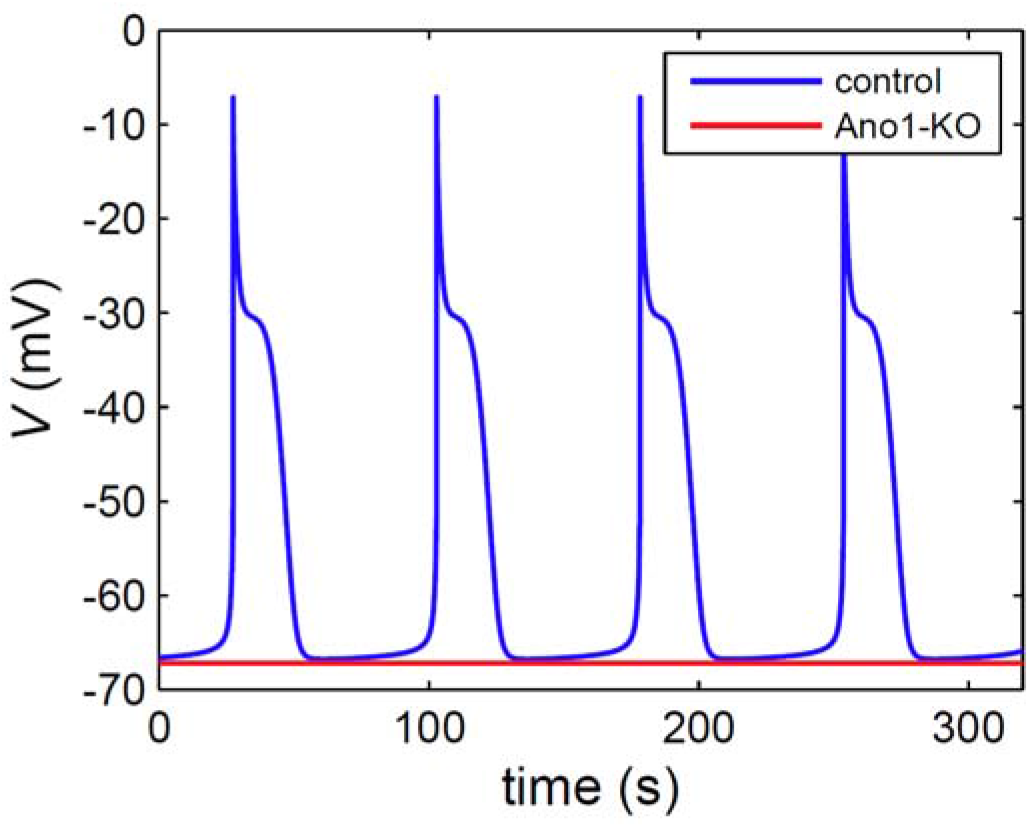
The existing model [after that of Imtiaz et al. (2007)] produces membrane potential oscillations when Ano1 is included, but steady membrane potential with Ano1-KO.

To study the cause of the elimination of oscillations with Ano1-KO, we carried out phase-plane analysis of the model. Phase-plane analysis graphically shows the relationship between the different state variables, rather than plotting the variables against time. It is a common approach for analysing models of voltage or calcium oscillations (Keener & Sneyd 2009). It can reveal the underlying mechanistic structure which causes the waveforms to have the shape they do. To analyse the cause of oscillations and the coupling between the M-clock and the C-clock, we separated the phase-plane analysis for the two clocks and used a novel approach to represent the coupling between the clocks graphically.

Figure 4(a) shows the phase plane for the C-clock of the existing model, for two constant values of the L-type calcium-channel current. The figure also shows the nullclines of the system for the two values of current. A nullcline is the locus of all points where one variable does not change with respect to time, e.g., the *Y*-nullcline, where *dY/dt* = 0. The points where nullclines intersect are equilibrium points for the system i.e., both *Y* and *Z* are unchanging. Here, the *Y*-nullcline does not vary with different values of *I*_L_, whereas the *Z*-nullcline does. The plot shows superimposed trajectories for the two currents, in each case starting from the arbitrary initial point *Z* = 1.5 μM, *Y* = 0. For *I*_L_ = 0, where the two nullclines intersect on the negative-slope region of each, there is no possible steady state. There is an equilibrium point at the intersection, but it is unstable. The trajectory settles to a periodic clockwise-rotating orbit, where each cycle retraces the preceding one, around this unstable equilibrium point. In contrast, for *I*_L_ = −10 pA, both the *Y*- and *Z*-nullclines have positive slope at their intersection, which represents a stable equilibrium point at a high calcium concentration, *Z* = 3.6 μM. The trajectory from the same initial point moves towards the two nullclines, overshooting the *Z*-nullcline then settling at this steady state. The high Ca^2+^ concentration is consistent with the higher influx of calcium through the L-type channel. These constant-*I*_L_ situations show the possible actions of the C-clock when it is decoupled from the M-clock. Decoupling is possible as the L-type calcium channel is the only mechanism in the model for the M-clock to modify calcium concentrations directly. The decoupled representation shows that the C-clock can oscillate without requiring the M-clock, and that the M-clock can eliminate these oscillations by providing sufficient calcium influx via the L-type channel.

**Figure 4.**
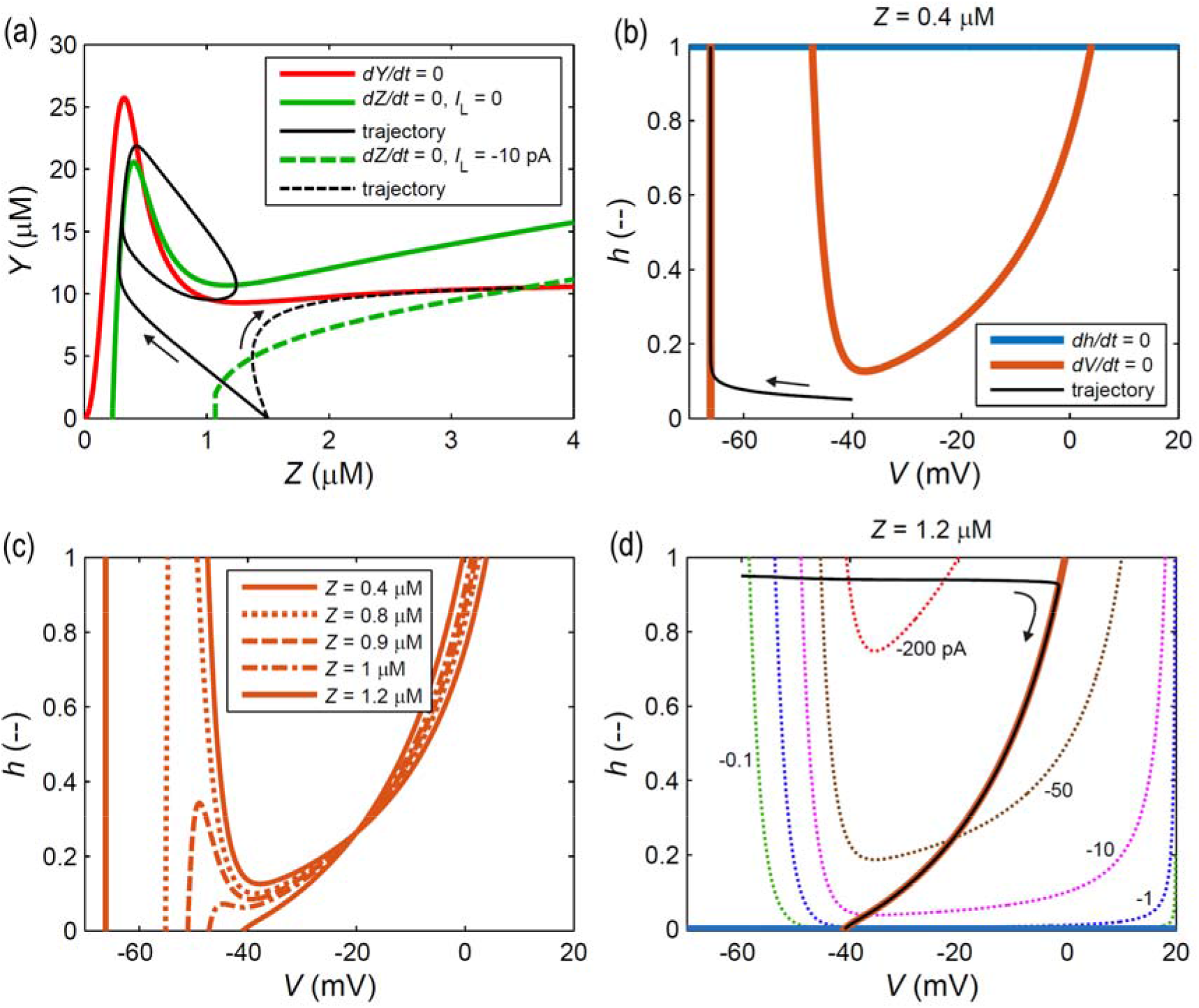
C-clock and M-clock phase planes for the original model. *h* = inactivation gate state, *V* = membrane potential, *Y* = store [Ca^2+^], *Z* = cytosol [Ca^2+^]. (a): nullclines and simulations for the C-clock, for two values of L-type current. (b): nullclines and a trajectory of the M-clock for intracellular calcium concentration 0.4 μM. (c): *V*-nullclines for several values of *Z*. (d): nullclines and a trajectory of the M-clock for calcium concentration 1.2 μM, superimposed on contours of constant L-type channel current in pA.

Figures 4(b) shows a similar phase-plane analysis for the M-clock, decoupled from the C-clock. The decoupling is achieved by plotting the phase plane for a constant value of the intracellular calcium concentration *Z*: 0.4 μM in Figure 4(b). This value approximates the minimum of the cyclic variation in *Z* shown in Figure 4(a). Since in the existing model *h*_∞_ is independent of *V*, the *h*-nullcline is always a horizontal line, which, for the value of *Z* shown in Figure 4(b), is at *h* = 1. The *V*-nullcline is a cubic curve having a single (unattainable) maximum at *h* >> 1, somewhere near *V* = −58 mV, and a minimum at *h* = 0.13. Nullclines of positive slope attract; thus, the left-hand arm of the *V*-nullcline divides the phase plane into a region to its left where *dV/dt* > 0 and one to the right where *dV/dt* < 0. Nullclines of negative slope repel, so a starting point to the right of the middle arm would be attracted to the right-hand arm. There are three equilibrium points (where the *V*-nullcline intersects *h* = 1), of which the outer two are stable and the middle one is unstable. A trajectory starting at *V*= −40mV, *h* = 0.05, is in a region where *dV/dt* << 0 and *dh/dt* > 0, so it tracks rapidly to the left, reaching the left-hand arm of the *V*-nullcline in about 0.15 s. Being still in a region where *dh/dt* > 0, it then climbs the *V*-nullcline at a rate controlled by *τ_h_*, reaching the stable equilibrium point at *V* = −68 mV, *h* = 1, in about 8 s.

As shown in Figure 4(c), the shape of the *V*-nullcline is highly *Z*-dependent, because *Z* affects the Ano1 chloride-ion current, as seen in eq. 9. The turning point (maximum) of the *V*-nullcline progressively reduces in height as Z increases. If *Z* reaches the value of 1.2 μM, there is no longer any reversal in the *V*-nullcline, at least in the attainable region 0 ≤ *h* ≤ 1, and the situation is as shown in Figure 4(d). This value of *Z* approximates the maximum of the cyclic variation in *Z* from Figure 4(a).

*Z* affects the M-clock also via the steady-state L-type inactivation gate *h_∞_*, as seen in eq.12. With Z = 1.2 μM, the *h*-nullcline is at *h* = 0. The available part of the *V*-nullcline is now a single curve of positive gradient. There is a stable equilibrium point near *V* = −40mV, *h* = 0, and a trajectory starting at *V* = −60 mV, *h* = 0.95, approaches this equilibrium point via a path which has its maximum *V*-value where it first reaches the *V*-nullcline. The path taken shows that the attraction to the *V*-nullcline is much stronger than that to the *h*-nullcline, i.e., at the starting point, *dV/dt* >> 0 whereas *dh/dt* is only slightly less than zero, being again limited by the time constant *τ_h_*.

Together, Figures 4(b and d) approximately define what happens during an action potential (AP). In Figure 4(d), starting with *h* and *Z* both high, the trajectory represents the upstroke and declining plateau of the action potential. Initially *dV/dt* >> 0 because the starting point is a long way from the *V*-nullcline, and there is a rapid increase in voltage until the trajectory reaches the *V*-nullcline, then, with the operating point under the weaker attracting influence of the *h*-nullcline at *h* = 0, the path gradually descends the *V*-nullcline. If it be now imagined that the simultaneous traversing of the (*Z*, *Y*)-orbit has reduced *Z* to values of the order of 0.4, the picture changes to that shown in Figure 4(b). The trajectory there effectively continues that shown in Figure 4(d), representing first the rapid final falling voltage of the downstroke at the end of the AP, then the very slow and slight rebuilding of voltage during diastole, wherein the trajectory approaches the *h*-nullcline which has now removed to *h* = 1. Changes in calcium concentration *Z* affect both the *h*- and *V*-nullclines. During diastole, *Z* increases, reducing and ultimately abolishing the left arm of the *V*-nullcline in (*V*, *h*)-space. As the peak of the *V*-nullcline falls below the value of *h* to which the operating point has climbed, the next action potential is triggered. This same increase in *Z* also brings down the *h*-nullcline, inactivating the L-type calcium channel.

Figure 4(d) also shows contours of constant L-type channel current in (*V*, *h*)-space, thus defining how the M-clock acts as an input to the C-clock. During the AP upstroke, i.e., the almost horizontal part of the trajectory in Figure 4(c), the current is large and passes through a maximum, whereas it is small at low and increasing *h* with low *V*, prior to the AP trigger.

Together, the plots in Figure 4 show that the C-clock drives the oscillations in the M-clock, and show the sequence and coupling of the two clocks over a full oscillation. The sequence starts with low calcium concentration *Z* and low L-type current *I*_L_. The calcium oscillations represented in Figure 4(a) increase *Z* enough to cause the M-clock to move from the state in Figure 4(b) towards that shown in Figure 4(d). This change triggers the action potential through the change in the *V*-nullcline as described above, causing a large inrush of calcium ions as L-type current peaks. This can be seen in Figure 4(d), where the horizontal part of the trajectory passes through a region where *I*_L_ exceeds 200 pA in magnitude. The high L-type current temporarily abolishes the Z-nullcline peak, causing the calcium trajectory to divert from its orbit towards a high-*Z* equilibrium point near *I*_L_ = −10 pA (Figure 4a), completing the transition of the M-clock from the state shown in Figure 4(b) to that in Figure 4(d). As the action potential ends, the now-small value of the gating variable *h* inactivates the L-type calcium channel. The L-type inactivation allows a reduction of *Z* in the C-clock, allowing the M-clock to return to its initial state in Figure 4(b). The oscillations of the C-clock then initiate the process again. The entire sequence of these events for the recombined and interacting M- and C-clocks is shown in cine-images in the Supplement.

Figure 5 shows the effect of Ano1-KO on the M-clock phase plane. Removing Ano1 has no effect on the *h*-nullcline, which continues to move up and down with the value of the calcium concentration *Z*, but the *V*-nullcline becomes fixed, i.e., independent of *Z*, because *Z* can no longer affect *V* directly; see eq. 3. This invariant *V*-nullcline removes the possibility of triggering an AP, because trajectories approach and remain at the left arm of the *V*-nullcline. The only exception to this occurs when *Z* is small enough that the *h*-nullcline intersects all three arms of the *V*-nullcline. For instance, if *Z* = 0.6 μM (not shown), the *h*-nullcline is at *h* = 0.93. For such smaller values of *Z* there are two stable equilibrium points where the *h*-nullcline intersects the left and right arms; depending on start position, trajectories will end at one of these two points.

**Figure 5.**
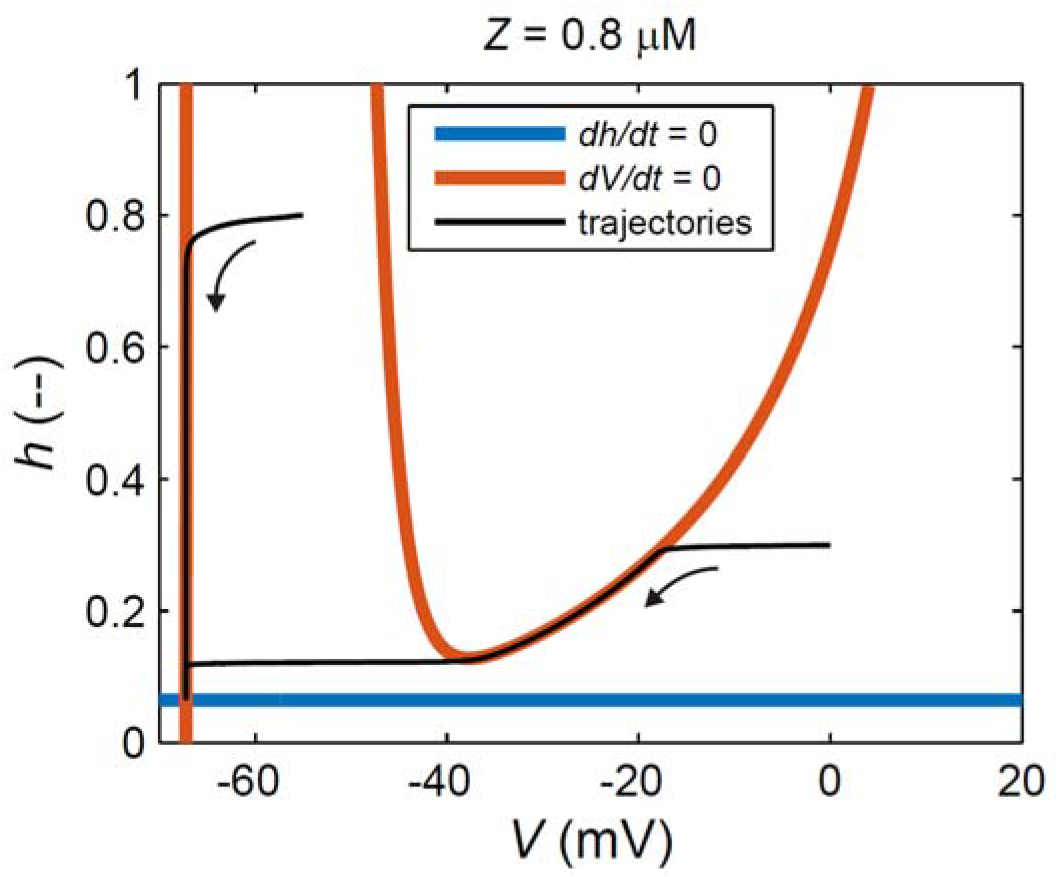
M-clock phase plane for the original model with Ano1-KO and intracellular calcium concentration *Z* fixed at 0.8 μM. Trajectories starting at any (*V*, *h*) location eventually end up at *V* = −67 mV, *h* = 0.08.

Figures 4 and 5 demonstrate that the M-clock cannot sustain oscillations by itself, i.e., in the absence of the C-to-M coupling afforded by the Ano1 channel, as it cannot oscillate without there being a varying calcium concentration. Therefore, the model requires modification if it is to have the capacity to match the Ano1-KO data shown in Figure 1.

### 3.2 Analysis of the new model

We now modify the model to make the inactivation gating variable for the L-type calcium channel depend on voltage *V* as well as calcium concentration *Z*, by setting *p* = 1 in eq. 12 and *τ*_x_ > *τ*_n_ in eq. 13. The *h*-nullcline in Figure 6 (left) shows the form of the *h*_∞_(*V*)-dependence so created; a similar curve defines the *V*-dependence of the time constant with *τ*_x_ and *τ*_n_ the maximum and minimum values respectively. The values of some other parameters are also changed, as shown in the right-hand column of Table 1.

**Figure 6.**
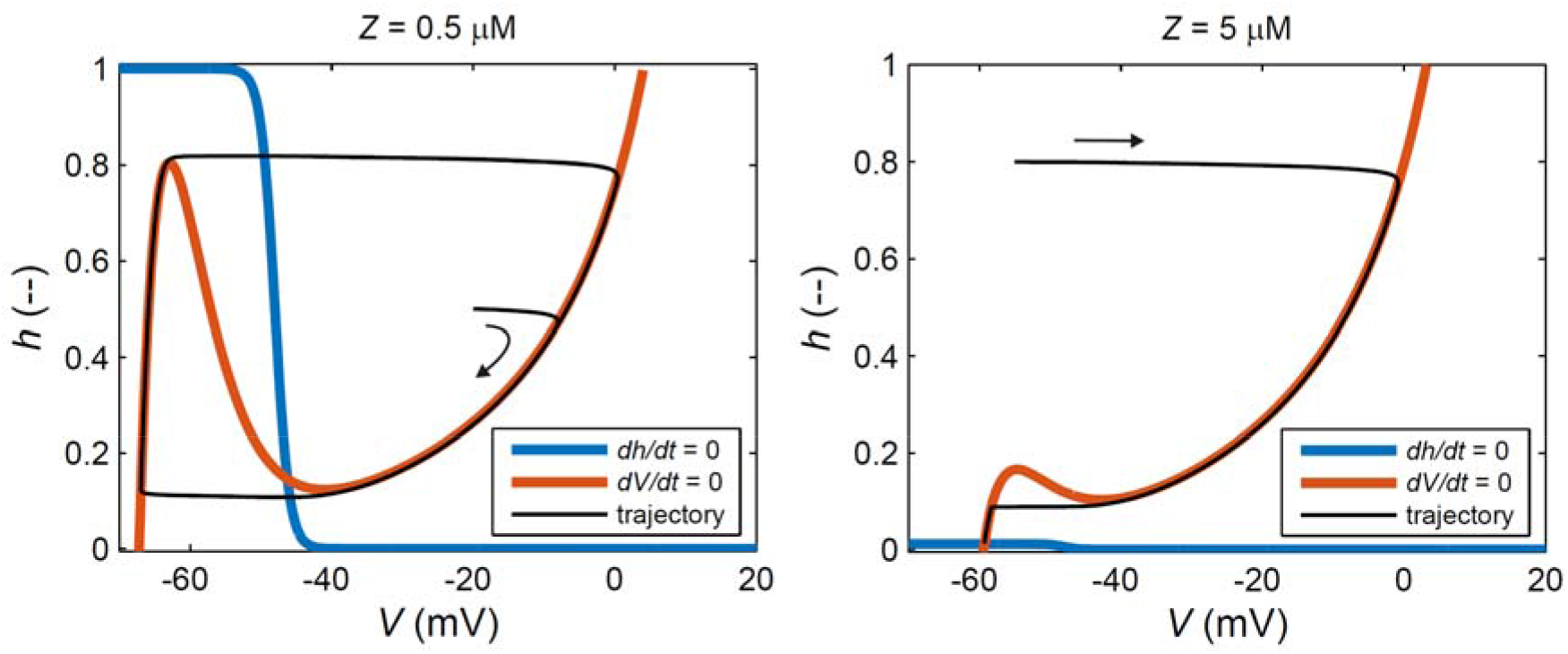
The M-clock phase plane for the new model. Left: oscillations for low *Z*. Right: no oscillations with high *Z*.

Figure 6 (left) shows the modified M-clock phase plane, for a low calcium concentration, *Z* = 0.5 μM. The *h*-nullcline is sigmoidal with respect to voltage, while the *V*-nullcline remains qualitatively as it was in Figure 4, although, because of the various parameter-value changes, its position relative to *Z*-value has changed compared to Figures 4 and 5. The modified *h*-nullcline intersects the *V*-nullcline at a single point where the *V*-nullcline has negative gradient, meaning that this equilibrium point is unstable and resides inside a clockwise-orbiting cyclic trajectory. The trajectory follows the two external arms of the *V*-nullcline, with a jump between those arms once each turning point has been passed. The jump from the left to the right arm of the *V*-nullcline is the AP upstroke, while the decrease of *h* along the right arm constitutes the declining AP plateau, and the jump from the right to the left arm constitute the rapid downstroke at the end of the plateau phase. When the calcium concentration *Z* is sufficiently high, the situation is as in Figure 6 (right). Despite the unchanged *h*(*V*)-relation, the *h*-nullcline is now close to *h* = 0 for all *V* because *h*(*Z*) so decrees, and the phase plane shows that oscillations are now impossible, since the two nullclines now intersect at a stable equilibrium point to which all trajectories are eventually bound.

Simulated membrane potentials from the new model, with and without Ano1, are given in Figure 7(a). The membrane potential oscillates under both control (experimentally, wildtype) and Ano1-KO conditions. The oscillations for Ano1-KO (*G*_xC_ = 0, eq. 9) have a lower frequency, larger maximum, and lower minimum than the oscillations for wild-type conditions. These behaviours are qualitatively consistent with the experimental data in Figure 1, and, because of the several changes in parameter values detailed in the right-hand column of Table 1, the frequencies of both control and Ano1-KO waveforms are not dissimilar to those shown in Figure 1. However, the AP duration increases slightly under Ano1-KO, which is a point of difference, and the model does not produce the considerable change in AP morphology between wild-type and Ano1-KO experiments.

**Figure 7.**
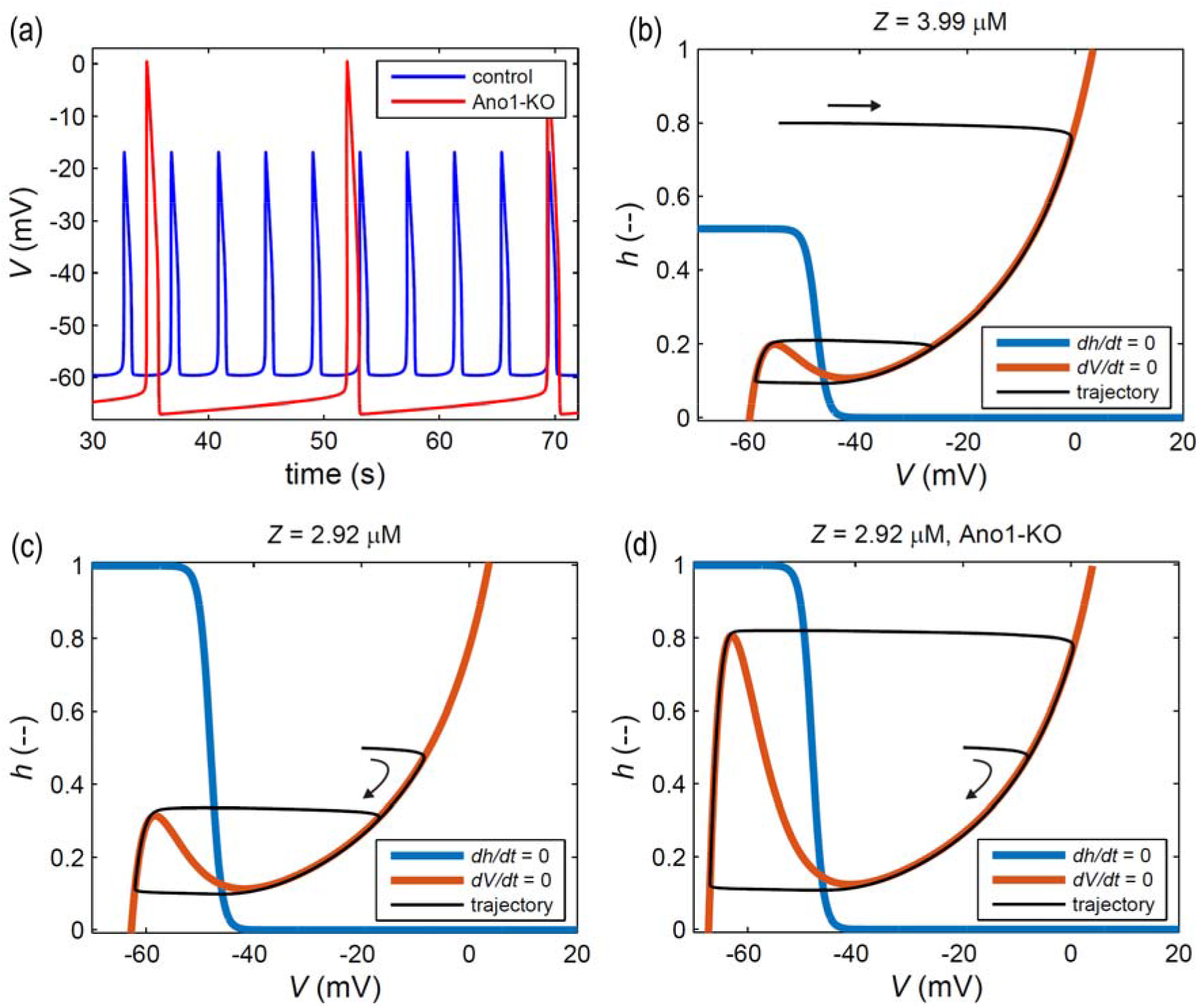
The effect of Ano1 on simulated membrane potential in the new model. (a): *V*(*t*)-traces from the full new model under (blue) control (AP frequency 14.7 /min) and (red) Ano1-KO (frequency 3.46 /min) conditions. (b): the phase plane of the isolated M-clock from the new model when the value of *Z* is the maximum of the cyclic variation in the full model under control conditions. (c): as (b), but when *Z* is the minimum of the cyclic variation in the full model. (d): as (c), but under Ano1-KO conditions. The *V*-nullcline and the trajectory would be unchanged if *Z* had the value shown in (b).

Figure 7(b–d) shows the corresponding M-clock phase plane. In panels (b) and (c) are shown the extremes of the variation of the *V*- and *h*-nullclines under control conditions; the values Z = 3.99 and 2.92 μM are the maximum and minimum of the cyclic variation of *Z*(*t*) in the full model under control conditions. Throughout the cycle of control oscillation in the full model (panel a), the turning point of the *V*-nullcline’s left arm occurs at a much lower value of *h* (and a slightly higher value of *V*) than is the case when Ano1 current is absent, as in panel (d). Consequently, the peak-to-peak AP voltage excursion is smaller in control conditions, as there is a smaller distance between the left and right arms of the *V*-nullcline at the height (*h*-value) where the AP occurs. This smaller excursion is made up of both a higher starting point and a lower summit, for reasons that are again obvious from comparison of the three phase planes. In addition, with the turning point located further away from the *h*-nullcline in control conditions, the operating point does not traverse the region closer to the *h*-nullcline where *dh/dt* is smaller and the rate of progression correspondingly less. Thus, because Ano1 current triggers the AP at a lower value of *h*, normal Ano1 operation results in a shorter diastolic interval between action potentials. The entire sequence of interacting M- and C-clock phase-plane events, for the traces in Figure 7(a) from the full new model, is shown in cine-images in the Supplement.

## 4 DISCUSSION

Imtiaz et al. (2007) declared their model to be “qualitative, … encapsulat[ing] the essential features of the experimental data and interrelationships of observed variables”, even though explicit quantitative values of parameters were given. We have attempted to reinterpret their model in as close to a fully quantitative manner as possible. This required some parameters to be substituted altogether. Thus, what they gave as a time constant *τ*_m_ is in our model the capacitance, and conductances which they gave in units of mS we use as nS. A further difficulty concerns the constant *α* defined by the laws of physics as in eq. 8; the value so given is orders of magnitude at odds with the non-dimensioned value in their paper. We also include a constant *b*_c_ in eqs. 1 and 2 defining the proportion of Ca^2+^ ions which are bound by buffers in the cytosol and in the store. This constant, which was omitted from their model, effectively slows the time-base of the calcium model by a factor of 10^2^. As a result, the control or ‘wild-type’ frequency in Figure 3 is only 0.795 /min, whereas Figure 1 shows a corresponding frequency of 20.9 /min. Furthermore, the simulated potential in Figure 3 reaches a minimum of −66.8 mV, whereas the experimental one in Figure 1 does not go below −46 mV. We ascribe the large difference in frequency mainly to this source, noting however that *b*_c_ has yet to be measured in lymphatic muscle cells. In principle, a mathematical model can be rescaled to satisfy any given time-base, but in practice such freedom is not available in a model intended to approximate realistic values of parameters, because many of these are constrained by experiment.

Nevertheless, even this relatively simple model (by the standards of most recent models of electrochemical pace-making) has many parameters, and so the total parameter-value space is large. Consequently, for the new model we have been able to find parameter values that permit a reasonable degree of approximation of the observed frequencies: 14.7 /min simulated vs. 20.9 /min observed under control conditions, 3.46 /min simulated vs. 5.11 /min observed for Ano1-KO. Furthermore, and more importantly, our new model successfully emulates the reduction in frequency by a factor of about four caused by Ano1 knock-out experimentally (4.25 simulated, 4.09 observed), whereas the existing model did not produce action potentials at all under these conditions.

However, there is a trade-off; whereas the existing model emulated rather well the morphology of the control AP, with its pronounced initial spike and flat plateau (Figure 1), the APs from the new model display a chisel shape, i.e., they lack an initial spike, and the plateau slopes downward, in both control and Ano1-KO conditions (Figure 7a). Moreover, the diastolic minimum of the simulated control waveform in Figure 7a is −59.6 mV, an improvement over that shown in Figure 3 but still substantially below the observed value. The observed Ano1-KO AP is very short and has different morphology relative to the wild-type one, whereas the predicted one is slightly longer and of similar morphology. A further point of difference concerns the extent of gradual depolarisation during diastole. However, it should be noted that in further models (not described in detail here) where we have fully isolated the M-clock, i.e., calcium concentration *Z* is no longer present at all, we can emulate the Ano1-KO recording closely in almost all respects; see Figure 8. Accurately representing the AP shape in both wild-type and Ano1-KO conditions may require (e.g.) modelling of additional calcium and/or chloride microdomains; see Youm et al. (2019) for an example. Experimentally we also need a better understanding of the chloride equilibrium potential in lymphatic muscle cells.

**Figure 8.**
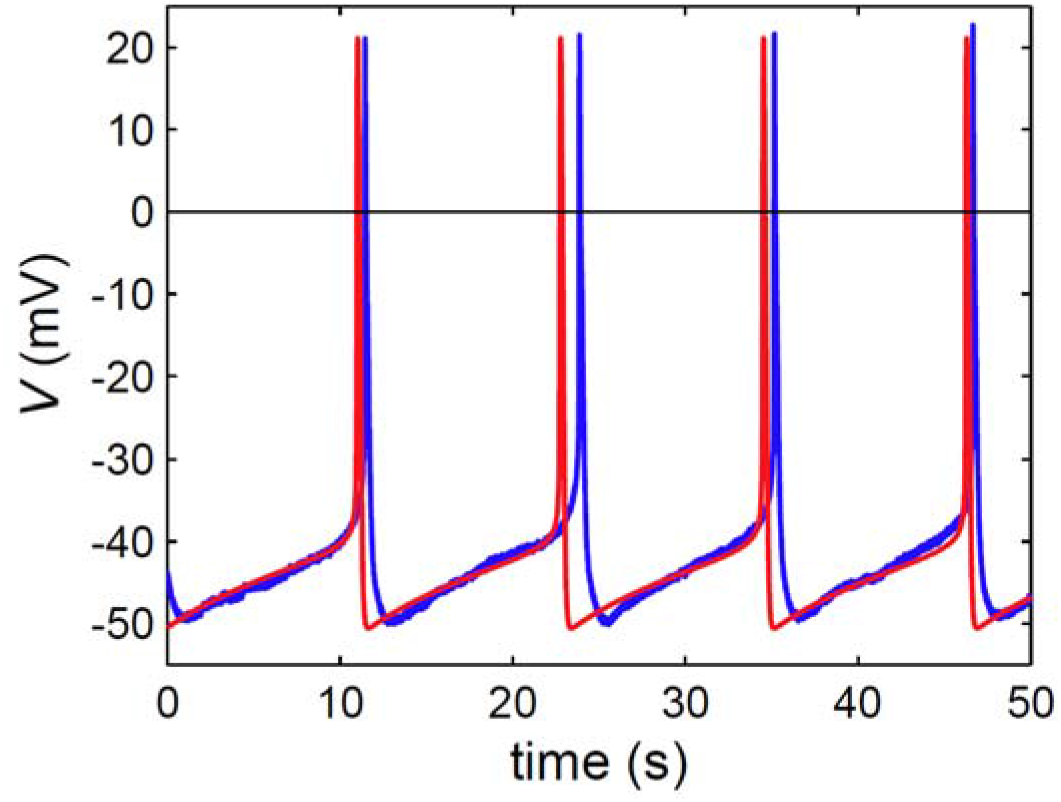
A comparison of the recorded Ano1-KO data (blue) with a trace from an M-clock-only model (red). The frequency of the simulated voltage is 5.10 /min; that of the recorded data varied slightly from beat to beat but averages 5.11 /min, based on the four peaks shown here.

In this paper we used simulations and phase-plane analysis to compare an existing and a new model of LMC pace-making oscillations. In developing this model of LMC pace-making, we have focussed on an older and simpler model, relative to the latest electrochemical models in most biological areas, containing just the core mechanisms required for oscillations of the C-clock and M-clock. We justify this approach by the need for a simpler model for phase plane analysis and the fact that very many recent models place the C-clock centrally (Dupont et al. 2016), regarding the oscillations of voltage as essentially an enslaved outcome of the C-clock oscillations. While cyclically varying calcium concentration is certainly the ultimate activator of rhythmic muscular contractions, this paradigm appeared unsuited to explaining the Ano1 knockout observations shown in Figure 1, whereby removing the mechanism which links cytosolic calcium ion concentration directly to transmembrane currents left action potentials intact, albeit at a much lower frequency and with greatly altered morphology.

Although our modification to the existing model is primarily focussed on a single mechanistic change—voltage dependence of the gating variable in the L-type calcium channel—any change in a relatively simple model such as is represented by eqs. 1–4 is important, and so the significant qualitative impact is not unexpected. The change made here results in the introduction of M-clock-driven oscillations and a corresponding increase in complexity in understanding the system. The model output differs from typical models of calcium oscillators in lymphatic and related cell types where the C-clock drives the M-clock. However, our changes are based upon the recent data (Figure 1), and the results show that the M-clock-driven oscillator can replicate the frequency reduction with Ano1-KO. Although the results rule out C-clock-driven oscillations, they are not inconsistent with the coupled-clock theory from cardiac pacemaker cells (Yaniv et al. 2015), as, although the C-clock is not essential for oscillations in the proposed model, it can help control the oscillatory frequency.

There is strong experimental support for the notion that L-type current inactivation is influenced by voltage as well as cytosolic calcium concentration, as described by Hille (2001). However, at least for cardiac pacemaker cells, the prevailing impression seems to be that voltage-dependent inactivation plays a lesser or minor role; see, e.g., Limpitikul et al. (2018). In contrast, in order to satisfy the observed behaviour of lymphatic muscle cells when Ano1 channels are knocked out pharmacologically or genetically, we believe that *V*-dependent inactivation of L-type current is essential. As we have demonstrated above, only thus can pace-making of reduced frequency be sustained after Ano1-KO, at least in the context of the admittedly simplified model with which we work. Voltage dependence for inactivation gating variables is also common in larger and more detailed mechanistic models of related cell types (e.g., Lees-Green et al. 2014).

In both the existing and novel models that we studied, Ano1 has the role of controlling the action potential initiation, but the consequences of this role differ considerably between the two models. In the existing model, rising calcium causes Ano1 to trigger the AP, which would not occur without this trigger. In contrast, increasing calcium in the novel model causes Ano1 to change the trigger point for the AP from a higher value of the gating variable *h* to a lower one. This change in the trigger point initiates the AP sooner, resulting in a higher frequency of oscillation. However, without Ano1, action potentials and oscillations still occur in the novel model.

To analyse the pacemaker oscillations, we introduced novel phase-plane analysis to represent the M-clock and the C-clock and their interactions graphically. In contrast, typical phase-plane analysis of pacemakers represents a single oscillator, rather than two coupled oscillators. This analysis allows us to understand complex interactions in the system and to separate the cause of the oscillations from their effect. The C-clock phase plane is similar to those for existing calcium oscillators (Dupont et al. 2016), as expected because the C-clock model is based on existing calcium models. In contrast, the M-clock phase plane (Figures 6 and 7) is analogous to the phase plane for the FitzHugh-Nagumo (FHN) model (FitzHugh 1961, Nagumo et al. 1962). This latter model is a two-dimensional reduction of the classical four-variable Hodgkin-Huxley model, wherein the potassium-channel gating variable is the equivalent variable to the L-type calcium-channel inactivation variable *h* here. The FHN model depends on a *V*-nullcline of cubic shape, with two turning points. In the original model, this was intersected by a straight line of positive slope, being the nullcline for the potassium-channel gating variable. Keener & Sneyd (2009) define the generalised FHN model, in which this line may be any suitable curve. Here, the phase plane is upside-down relative to the traditional portrayal: the *h*-nullcline has negative slope^1^, and the cubic curve has its local maximum to the left of its local minimum. As in the usual FHN model, here the voltage variable *V* has much faster temporal dynamics than the gating variable *h*, and so trajectories rapidly approach the *V*-nullcline, moving approximately horizontally, if starting far from it. The rapid approach to a *V*-nullcline represents the upstroke or downstroke of the action potential. Trajectories then track along the *V*-nullcline, with the direction and velocity dependent upon the dynamics of the gating variable *h*, as controlled through eq. 4 by the time constant *τ_h_*. A trajectory stops tracking the *V*-nullcline either where it reaches a turning point in the *V*-nullcline or when the C-clock has changed the location of the *V*-nullcline, both resulting in the trajectory rapidly moving to a new arm of the *V*-nullcline. Together, the *V*- and *h*-nullclines allow one to observe the qualitative behaviour of the system and the shape of the action potential on the M-clock phase plane.

We have also introduced a novel graphical representation of the coupling between the two clocks. By comparing M-clock phase planes for different values of calcium concentration *Z*, we showed how the M-clock varies as a function of the C-clock trajectory. The graphical representation of the coupling shows how the oscillating trajectory (limit cycle) in the C-clock is abolished by sufficiently large L-type current. We have also used the L-type channel current to represent the coupling in the other direction; it is the primary means whereby the M-clock modifies the C-clock, and it effectively reduces the coupling from a function of two variables (i.e., *Z* and *h*) to just one (i.e. *I_L_*). We have superimposed C-clock phase planes with different L-type channel currents to show the effect of *I_L_*, while also plotting *I_L_* isoclines on the M-clock. Together, these additions explicitly show how the C-clock phase plane changes as a function of the M-clock trajectory. In particular, they show how the value of *Z* changes the location of the equilibrium in the M-clock in the original model, and how large values of *Z* remove the oscillating trajectory in the M-clock of the new model. The *I_L_* isoclines also show in a phase-plane representation that the L-type current stays at a relatively low level until the action potential is initiated, at which point it increases drastically. This enables one to observe all stages of the periodic sequence on the phase plane, particularly the all-important action potential trigger.

By starting from a relatively simple model consisting of only four ordinary differential equations, we have been able to deploy phase-plane analysis. Even with this simplification, the approach of representing the pacemaker via two phase planes is imperfect as the four-dimensional model is fully coupled and does not comprise two separate two-dimensional models. However, our approach provides valuable insights into the two oscillators and the coupling between the two. If in future we move to more mechanistically detailed models, complications could occur in using the approach, since approximations and additional assumptions will be needed to reduce such models to a form which is still amenable to this style of analysis.

While our focus in this paper has been on phase-plane analysis of LMC models, an important future direction of research will be parameter estimation and model validation of the novel model from experimental data. We have taken a first step in this direction by fitting a further developed M-clock model derived from that of eqs. 3 and 4 to the LMC Ano1-KO voltage data from Figure 1. In Figure 8, the model and data can be observed to match closely, providing evidence for both the M-clock driving of oscillations and the need for voltage dependence of the gating variable *h* in the models. Future model fitting will be required for the full model to simulate both wild-type and Ano1-KO data successfully. While in this paper our focus was on modifications to the M-clock stemming from the phase plane analysis, future research may require modifications to the C-clock by incorporating recent developments in calcium models (Dupont et al. 2016, Sneyd et al. 2017). Refinements introducing other coupling mechanisms between the two clocks, such as dependence of *E*_L_ on intracellular calcium concentration *Z*, may also be required.

The alternative to our approach of developing a simpler model is to develop detailed mechanistic models involving large numbers of equations and their respective variables and parameters. There is a trade-off between these two approaches, where the detailed mechanistic model has the advantage of potentially being closer to reality and may provide a better understanding of specific mechanisms. However, it has the disadvantage of being more difficult to analyse and interpret. In our case this difficulty manifests itself in understanding the role of and the coupling between the M-clock and the C-clock. It can be difficult in a detailed model to determine the cause and effect between mechanisms and oscillatory behaviour. Detailed models also have many more parameters, the values of which can be difficult or impossible to determine by either fitting to data or obtaining from the literature. This will be an important direction for future research on the topic.

## 5 CONCLUSIONS

Modelling provides an important tool to help understand function and failure of oscillations in lymphatic muscle cells. Here, we have made progress towards this goal by extending both a model of these cells and its corresponding analysis methodology. We have modified a model by Imtiaz and colleagues to enable the M-clock to drive the C-clock oscillations and thus respond to challenge in a way consistent with experimental data. We have also provided phase-plane analysis to understand the underlying causes and interactions of these forms of oscillation. Both advances will be important in the ongoing attempt to identify the mechanisms causing lymphoedema and to find targets for its pharmacological treatment.

## ACKNOWLEDGEMENTS

This research, EJH, and in part CDB, were supported by NIH grant R01-HL-122578 to MJD. SDZ acknowledges NIH grant R00-HL-143198.

## COMPETING INTERESTS

The authors declare that they have no competing interests.

## CONTRIBUTIONS

The project was initially conceived by MJD and CDB. The model was developed by EJH, CDB and CM. SJZ performed the experiments, in the laboratory of MJD. EJH and CDB wrote most of the computer code, with contributions from CM. EJH, CDB and CM all participated in the analysis and interpretation of numerical data. SJZ and MJD conducted the analysis and interpretation of experimental data. EJH and CDB wrote the report, with review by CM. All authors reviewed and approved the manuscript prior to submission.

## Footnotes

1 This results from the definition of *h* in terms of activation (1 = open, 0 = closed) rather than inactivation (0 = open, 1 = closed).

## Notes

### Competing Interest Statement

The authors have declared no competing interest.

## REFERENCES

Adams KE, Rasmussen JC, Darne C, Tan I-C, Aldrich MB, Marshall MV, Fife CE, Maus EA, Smith LA, Guilloid R, Hoy S, Sevick-Muraca EM (2010) Direct evidence of lymphatic function improvement after advanced pneumatic compression device treatment of lymphedema. Biomedical Optics Express 1(1): 114–125

Baluk P, Fuxe J, Hashizume H, Romano T, Lashnits E, Butz S, Vestweber D, Corada M, Molendini C, Dejana E, McDonald DM (2007) Functionally specialized junctions between endothelial cells of lymphatic vessels. Journal of Experimental Medicine 204(10): 2340–2362. doi:10.1084/jem.20062596

Breslin JW, Yang Y, Scallan JP, Sweat RS, Adderley SP, Murfee WL (2019) Lymphatic vessel network structure and physiology. Comprehensive Physiology 9: 207–299. doi:10.1002/cphy.c180015

Davis MJ (2005) An improved, computer-based method to automatically track internal and external diameter of isolated microvessels. Microcirculation 12(4): 361–372. doi:10.1080/10739680590934772

Dupont G, Goldbeter A (1993) One-pool model for Ca^2+^ oscillations involving Ca^2+^ and inositol 1,4,5-trisphosphate as co-agonists for Ca^2+^ release. Cell Calcium 14(4): 311–322

Dupont G, Falcke M, Kirk V, Sneyd J (2016) Models of Calcium Signalling. Interdisciplinary Applied Mathematics, vol 43. Springer International Publishing, Switzerland. doi:10.1007/978-3-319-29647-0

Engeset A, Olszewski W, Jæger PM, Sokolowski J, Theodorsen L (1977) Twenty-four hour variation in flow and composition of leg lymph in normal men. Acta Physiologica Scandinavica 99(2): 140–148. doi:10.1111/j.1748-1716.1977.tb10364.x

FitzHugh R (1961) Impulses and physiological states in theoretical models of nerve membrane. Biophysical Journal 1(6): 445–466

Gashev AA (2008) Lymphatic vessels: pressure- and flow-dependent regulatory reactions. Annals of the New York Academy of Science 1131: 100–109. doi:10.1196/annals.1413.009

Hald BO, Castorena-Gonzalez JA, Zawieja SD, Gui P, Davis MJ (2018) Electrical communication in lymphangions. Biophysical Journal 115(5): 936–949. doi:10.1016/j.bpj.2018.07.033

Hille B (2001) Ion Channels of Excitable Membranes, 3rd edn. Sinauer Associates Inc., Sunderland MA

Keener J, Sneyd J (2009) Mathematical Physiology I: Cellular Physiology. Interdisciplinary Applied Mathematics, vol 8/I, 2nd. edn. Springer Science+Business Media LLC, New York. doi:10.1007/978-0-387-75847-3

Imtiaz MS, Zhao J, Hosaka K, von der Weid P-Y, Crowe MJ, van Helden DF (2007) Pacemaking through Ca^2+^ stores interacting as coupled oscillators via membrane depolarization. Biophysical Journal 92(11): 3843–3861. doi:10.1529/biophysj.106.095687

Lees-Green R, Gibbons SJ, Farrugia G, Sneyd J, Cheng LK (2014) Computational modeling of anoctamin 1 calcium-activated chloride channels as pacemaker channels in interstitial cells of Cajal. American Journal of Physiology – Gastrointestinal and Liver Physiology 306(8): G711–G727. doi:10.1152/ajpgi.00449.2013

Limpitikul WB, Greenstein JL, Yue DT, Dick IE, Winslow RL (2018) A bilobal model of Ca^2+^-dependent inactivation to probe the physiology of L-type Ca^2+^ channels. Journal of General Physiology 150(12): 1688–1701. doi:10.1085/jgp.201812115

Maltsev VA, Lakatta EG (2013) Numerical models based on a minimal set of sarcolemmal electrogenic proteins and an intracellular Ca^2+^ clock generate robust, flexible, and energy-efficient cardiac pacemaking. Journal of Molecular and Cellular Cardiology 59: 181–195. doi:10.1016/j.yjmcc.2013.03.004

McAllister RE, Noble D, Tsien RW (1975) Reconstruction of the electrical activity of cardiac Purkinje fibres. Journal of Physiology 251: 1–59.

Morris C, Lecar H (1981) Voltage oscillations in the barnacle giant muscle fiber. Biophysical Journal 35: 193–213

Mortimer PS, Rockson SG (2014) New developments in clinical aspects of lymphatic disease. Journal of Clinical Investigation 124(3): 915–921. doi:10.1172/JCI71608

Muthuchamy M, Gashev A, Boswell N, Dawson N, Zawieja D (2003) Molecular and functional analyses of the contractile apparatus in lymphatic muscle. FASEB Journal 17(8): 920–922. doi:10.1096/fj.02-0626fje

Nagumo J, Arimoto S, Yoshizawa S (1962) An active pulse transmission line simulating nerve axon. Proceedings of the IRE 50(10): 2061–2070

Noble D (1962) A modification of the Hodgkin-Huxley equations applicable to Purkinje fibre action and pacemaker potentials. Journal of Physiology 160(2): 317–352

Olszewski WL, Engeset A (1980) Intrinsic contractility of prenodal lymph vessels and lymph flow in human leg. American Journal of Physiology - Heart and Circulatory Physiology 239(6): H775–H783

Rockson SG, Rivera KK (2008) Estimating the population burden of lymphedema. Annals of the New York Academy of Science 1131: 147–154. doi:10.1196/annals.1413.014

Sanders KM (2019) Spontaneous electrical activity and rhythmicity in gastrointestinal smooth muscles. Advances in Experimental Medicine and Biology 1124: 3–46. doi:10.1007/978-981-13-5895-1_1

Scallan JP, Zawieja SD, Castorena-Gonzalez JA, Davis MJ (2016) Lymphatic pumping: mechanics, mechanisms and malfunction. Journal of Physiology 594(20): 5749–5768. doi:10.1113/JP272088

Seo Y, Lee HK, Park J, Jeon D, Jo S, Jo M, Namkung W (2016) Ani9, a novel potent small-molecule ANO1 inhibitor with negligible effect on ANO2. PLoS One 11(5): e0155771-1–e0155771-16. doi:10.1371/journal.pone.0155771

Sneyd J, Han JM, Wang L, Chen J, Yang X, Tanimura A, Sanderson MJ, Kirk V, Yule DI (2017) On the dynamical structure of calcium oscillations. Proceedings of the National Academy of Sciences of the United States of America 114(7): 1456–1461. doi:10.1073/pnas.1614613114

To KHT, Gui P, Li M, Zawieja SD, Castorena-Gonzalez JA, Davis MJ (2020) T-type, but not L-type, voltage-gated calcium channels are dispensable for lymphatic pacemaking and spontaneous contractions. Scientific Reports 10: 70-1–70-24. doi:10.1038/s41598-019-56953-3

Trzewik J, Mallipattu SK, Artmann GM, Delano FA, Schmid-Schönbein GW (2001) Evidence for a second valve system in lymphatics: endothelial microvalves. FASEB Journal 15(10): 1711–1717. doi:10.1096/fj.01-0067com

Yaniv Y, Lakatta EG, Maltsev VA (2015) From two competing oscillators to one coupled-clock pacemaker cell system. Frontiers in Physiology 6: 28-1–28-8. doi:10.3389/fphys.2015.00028

Youm JB, Zheng H, Koh SD, Sanders KM (2019) Na-K-2Cl cotransporter and store-operated Ca^2+^ entry in pacemaking by Interstitial Cells of Cajal. Biophysical Journal 117(4): 767–779. doi:10.1016/j.bpj.2019.07.020

Zawieja SD, Castorena JA, Gui P, Li M, Bulley SA, Jaggar JH, Rock JR, Davis MJ (2019) Ano1 mediates pressure-sensitive contraction frequency changes in mouse lymphatic collecting vessels. Journal of General Physiology 151(4): 532–554. doi:10.1085/jgp.201812294

